# Non-detection during excursions by citizen scientists modeled as a function of weather, season, list length, and individual preferences

**DOI:** 10.1101/2024.09.30.615418

**Authors:** Gert W. Jacobusse, Eelke Jongejans

## Abstract

**INTRODUCTION:** Citizen science is an increasingly valuable source of information about biodiversity. It is challenging to use this information for analysis of distribution and trends. The lack of a protocol leads to bias in observations and therefore data are not representative. The bias is a consequence of unequal detection probabilities, caused by different preferences and habits of citizen scientists.

**METHODS:** We propose to incorporate characteristics of these excursions in analyses of data collected by citizen scientists to improve estimates of the probability that a species is not detected and reported, even though it does occur. By limiting these models to areas that are known to be occupied, detection can be modeled separately without considering variation in occupancy. We apply this idea to 150 common species in the Southwest Delta of The Netherlands, and illustrate the data selection, the modeling process and the results using four species.

**RESULTS:** The strongest features to predict detection are the number of species during a visit (list length), earlier observations of the target species by the same observer, and the day of year. We compare three approaches to predict the total non-detection probability that takes all visits to an area into account. Predictions based on only the number of visits were outperformed by predictions that also take the list length into account. Our predictions based on all features combined consistently beat both other approaches, across all 10 species groups that were compared.

**DISCUSSION:** We thus show that explicitly modelling the characteristics of all visits to an occupied area results in estimation of non-detection probabilities, while providing insight into the causes of detection and reporting bias. Furthermore, predictions of our model provide a basis for quantifying the sampling effort in each area, which is a promising first step to correct bias in citizen science data when aiming to map a species’ distribution.

## INTRODUCTION

Biodiversity plays a fundamental role in ecosystem functioning and resilience (Isbell et al.; 2015, Oliver et al., 2015). Numerous threats such as habitat loss, disturbance, pollution and climate change contribute to the decline of species. To address these threats effectively, it is essential to establish a quantitative understanding of biodiversity across regions (Hochkirch et al., 2021). Data gathered through citizen science are recognized as a valuable source of information about biodiversity (van Strien et al., 2013). Online platforms and the availability of technical aids like automatic species recognition from photos have made it easier to report observations during excursions. These developments increased the amount and quality of available data (Luna et al., 2018). However, the non-systematic nature of observations reported by citizen scientists causes the bias in the resulting datasets to be extensive. Most challenging is the lack of explicit information about the absence of species, as typically only species presences are reported. Biased detection and reporting need to be addressed first, to unlock the full potential of citizen science as a means for quantifying biodiversity. Researchers have recognized and investigated the consequences of bias in previous studies (Isaac et al., 2014; Ranc et al., 2017; Jha et al., 2022), but adequate tools to account for detection and reporting biases are dearly missing.

### Species distribution models

For the quantification of biodiversity across regions, a major consideration is the occupancy of areas by species. Species Distribution Models (SDM) predict where species live (occupied areas), based on climatic and environmental data (Elith & Leathwick, 2009; Melo-Merino et al., 2020). Ideally, SDMs are fitted to observations of both presences and absences of a species, where documented absences can come from both unoccupied and occupied areas (left part of Figure 1). In practice, however, many data sources, especially those gathered through citizen science, only contain presence data (Johnston et al., 2023). These presence data are typically incomplete because not all areas have been systematically searched or even visited (middle part of Figure 1). When presence records are not representative because of unequal detection probabilities, and species are not detected in some areas that they do occupy, this may lead to serious bias in parameter estimates in SDMs (Gu & Swihart, 2004). Unequal probabilities of a species being detected at least once in an area, are affected by multiple factors, including variable sampling efforts, which is usually thought of as the number of visits to an area. Here we focus on the opposite: the local probability that a species is not detected and reported during any of the visits in a given year, while the area is known to be occupied (based on evidence from previous years).

**Figure 1.**
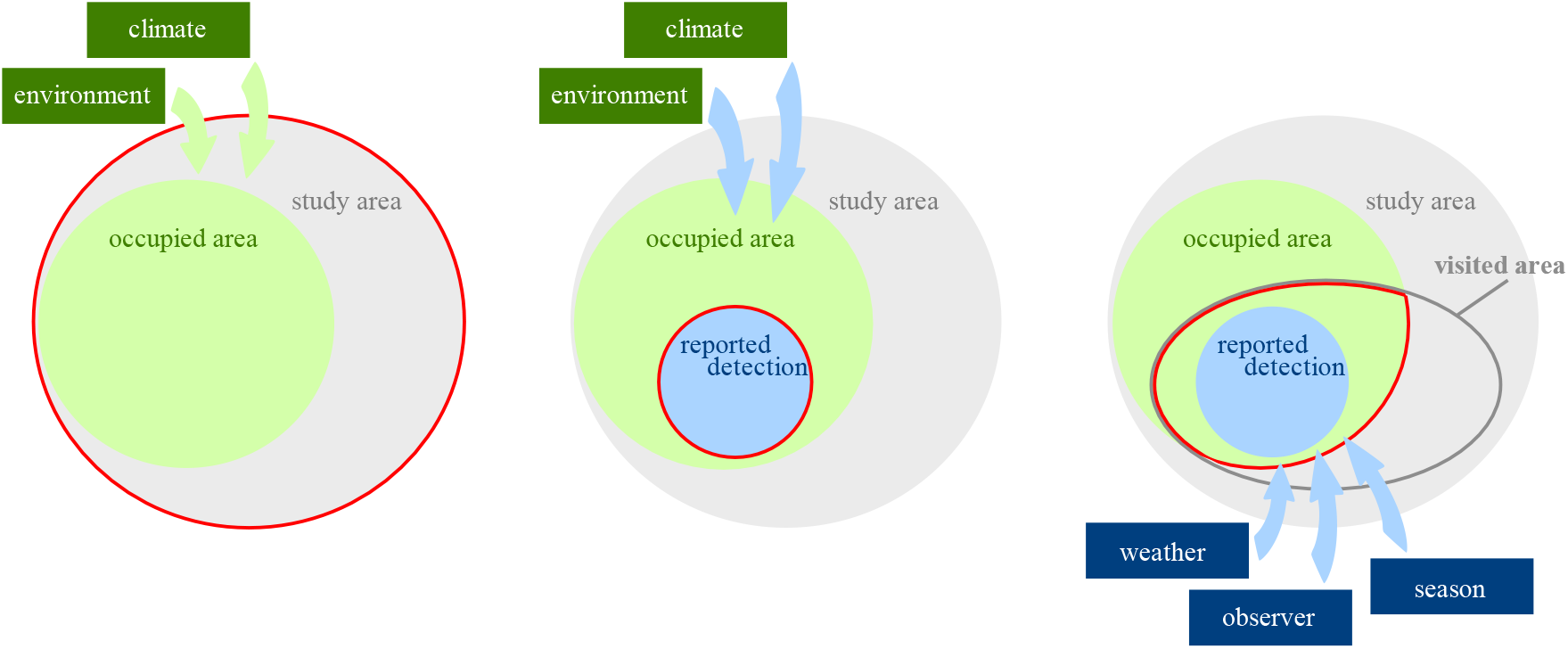
Schematic representation of a study area (grey) that is partly occupied by a species (green). Red borders indicate what data are available for the three modeling approaches. Left: ideal situation for fitting a Species Distribution Model with complete cover of the entire study area, resulting in both presence and absence data. Here, the SDM is a function of climate and environmental variables and can generate predictions for areas outside the study area. Mid: often, only presence data are available (blue), only for a subsection of the occupied area. SDMs require assumptions and generation of pseudo-absences. Right: In this manuscript, we focus on the modeling of non-detection probabilities, for which we focus on visits by citizen scientists to areas that are known to be occupied. Data therefore consist of both detections and non-detections that can be related to the frequency and seasonal timing of visits, and the characteristics of the visits like weather and observer characteristics.

### Modeling non-detection

Occupancy models distinguish between occupancy as the ecological quantity of interest and detection as a consequence of the observation process. They exploit replicate samples in each area to estimate the detection probability, assuming that the occupancy status of the area does not change (Royle & Dorazio, 2008). Many occupancy models incorporate time-varying and site-specific explanatory variables to account for unequal detection probabilities (MacKenzie et al., 2002; Tyre et al., 2003; Bailey et al., 2014). Some authors pointed out that careful sampling design and a strict definition of occupancy are necessary to achieve valid results (Kendall & White, 2009; Efford & Dawson, 2012) and provide insights into how to prioritize efforts to systematically gather data (Sanderlin et al., 2019). Given the lack of any strict definitions for citizen science data, this study will focus on opportunities to achieve valuable insights from more messy data. This is far from trivial, because even when data are gathered systematically, modeling non-detection is a challenging task.

The fact that observations are a consequence of both occupancy and detection introduces confounding, together with a lot of free parameters and therefore a need to limit complexity of the model. Research on non-detection has accomplished this by limiting the number of covariates and fitting quite restricted linear models, fixing detection probabilities per species (Kéry and Royle, 2008) or per visit (Outhwaite et al., 2018), applying special estimation procedures (Lele et al., 2012) and filtering species that show low detection probabilities (Ferrer-Paris & Sánchez-Mercado, 2020).

### Detection and occupancy

In the context of citizen science data, detection probabilities are usually very low, well below 10 percent. Typically, detection is not the result of a systematic effort but depends on the focus and preference of the observer, who is free to record, or not record, any of the species observed. Detection takes place only if the species is present in the area, observed during a visit, recognized, and reported. Each citizen scientist may have their own definition of an area, therefore areas for modeling need to be constructed in hindsight. Within these constructed areas, we define a visit list as the reported species (at least 1) in an area on a certain date by a certain observer. Defining a visit in this way is crucial because it means that the observer was present in the area on that date and had the chance to report other species that occupy the area. As input for modeling detection, we also define an excursion as the combination of all visits of an observer on one day, to one or multiple areas.

Occupancy also needs to be defined systematically. The incomplete overlap between home range and area, together with the unpredictable timing of visits, requires us to rely on cumulative observations of occupancy over time. Efford and Dawson (2012) referred to this as asymptotic occupancy, contrasting it with instantaneous occupancy, where each individual can only occupy a single area. Analogous to that, we will relax the assumption of closure – that there are no changes in occupancy between visits. In this study, we count an area as occupied during the entire year, even if the species is a migratory bird or an insect that can only be found during a specific part of the year. This relaxation carries some implications and risks that will be discussed later. The intended result is that presence of a species is not a necessary consequence of occupancy. Absence during the ‘wrong’ season can be accounted for by the model, with a zero detection probability. This is in line with the idea that allowing estimates of detection probability to vary between site visits has the potential to ‘absorb’ violations of the closure assumption (MacKenzie et al., 2017). On top of that, citizen scientists have a tendency to overreport interesting observations like a rare species, or a migratory bird that just arrived in the area. With the relaxed occupancy definition, this confounding between closure and detection can be accurately modeled as a seasonal increase in detection probability.

### Conditional models

Working with citizen science data and the aforementioned definitions of occupancy, visits and excursions makes modeling non-detection even harder, and increases the need to limit complexity. Therefore, we advocate species-specific models that are targeted at detection only, without considering occupancy. This can be done by conditioning analysis on areas that are known to be occupied. Like Chen et al. (2009) mentioned, inside areas that are known to be occupied, a zero observation is assumed to represent the overlooking of a species rather than its absence. Then non-detection can be directly modelled, without the need to account for possible non-occurrence by use of an occupancy model. This corresponds to the modelling approach in the right part of Figure 1. After the models are thus fitted using occupied areas only, the trained models will predict detection probabilities for all areas given the characteristics of the visits to those areas. Such predictions are also made for unoccupied areas: even completely unsuitable areas, like a land area for a fish, will receive a prediction. Therefore, it is unnecessary to a priori make a distinction between suitable and unsuitable areas.

In this paper we develop a model for species-specific non-detection in areas that are known to be occupied by that species, taking the frequency, timing and weather conditions of visits to an area into account as well as the list length and past reports of the observers. We test this non-detection model on citizen science data from the south-west part of the Netherlands. Apart from quantifying sampling efforts, this exercise will reveal species-specific insights into how explanatory variables affect detection probabilities at the visit level, which will allow researchers to map areas with insufficient sampling effort, which should eventually lead to improved species distribution modelling based on citizen science data.

## METHODS

### Modeling non-detection

We developed models to predict non-detection and tested these models using citizen science-based observation data from the Dutch National Database Flora and Fauna (NDFF), mostly from waarneming.nl, in the province of Zeeland from 2017 to 2023. Observations in 2023 are only used as test set. Observations before that are used to train the model and to determine which areas are occupied.

The unit of analysis is the unique combination of area, observer and date, which corresponds to a visit to an area during which at least one observation was done. By modeling per visit, specific effects of observer and weather variables can be expected to show up as important determinants of non-detection. In an approach with aggregated data per month or per area, these effects would be missed by averaging over longer periods or multiple observers. Areas have been defined as squared kilometers according to the Amersfoort / RD New (EPSG:28992) grid. All areas with at least 50 observations^1^ in total in the 2020-2022 period have been included.

A total of 150 species were selected from the 10 most observed groups, by taking the most common species within each group: 30 birds, 20 plants, 20 moths, 20 butterflies, 10 mammals, 10 dragonflies, 10 Diptera, 10 Hymenoptera, 10 Orthoptera and 10 mushrooms. For each species separately, a modeling dataset is created including only occupied areas: these are defined as areas where the species had been observed at least once in two out of three preceding years. For instance, observations in 2017-2019 determined which areas were known to be occupied (i.e., in at least two years) when using the 2020 data for training the model. Similarly, the 2021 and 2022 datasets were also used as training sets (see appendix 1 for an illustration). The training datasets form the basis to create a model that predicts the conditional probability of non-detection per visit in 2023.

The binary modeling target is whether or not the species has been observed during the visit. When a species is not observed, this is seen as a non-detection because only occupied areas have been selected for the modeling dataset. To predict the probability of non-detection, the following features are calculated to represent the circumstances during each visit:

- The list length, which is the number of species observed and reported during the visit
- The visit’s day of the year
- Temperature deviation on the day of the visit, compared to the average temperature on the same day of the year over the last thirty years, based on weather station Vlissingen
- For the observer, percentage of all observations before 2023 that were observations of the target species

Other features about weather (precipitation) and observer (average rarity of species observed, total number of observations, total number of species, overall and within the same group as the target species) have been calculated and compared, but were dropped during feature selection. Features about the observer only exist for observers that were already active before 2023, for new observers these features were encoded as -1, which allows the model to treat them as a separate category.

A RandomForestClassifier is trained to learn the relation between these features and the target. Using the trained model, non-detection probabilities are predicted for all visits in 2023. Cross validation is performed to create holdout predictions of probabilities for areas that did not take part in the training. This means that in the test set containing visits in 2023, each predicted probability for a visit is based on a model that has not used any visits to the same area during training. For validation, the Area Under Curve (AUC; Bradley, 1997) per visit is calculated to measure how well the model succeeds in predicting the non-detection of species in areas that are occupied. The AUC metric is a number between 0.0 and 1.0 that measures how well the ordering in the predictions matches the actual binary non-detection. The closer the value is to 1.0, the better the predictions match reality. Random ordering would result in an AUC of 0.5, so the expected range of modeling results is between 0.5 for a worthless model and 1.0 for a perfect model. For each species separately, the conditional non-detection probability P(non-detection|occupancy) per area is calculated as the product of the conditional non-detection probabilities of all visits to the area in 2023, assuming that the probabilities of different visits are independent. This prediction is tested against actual non-detection during all visits to that area combined, by calculating the AUC per area. The result is compared to two benchmarks for predicting detection: one based on only the number of visits and one that also takes the list length per visit into account.

Feature selection was an iterative process, involving both feature definition and feature selection. Guiding principles were: First, to avoid multicollinearity by standardizing features. For example, temperature has been expressed as a deviation and observations of the target species as a percentage. Second, features were selected in a forward fashion, starting with one feature per category (weather, observer and season), only adding more features when the improvement of the AUC per area was at least 0.01. For each selected feature, this improvement is reported as the contribution to the final model, calculated by running the model again without the feature to measure the difference in AUC per area, averaged over all species.

To create marginal dependence plots, the detection probability is calculated on a rolling window for each decile of the input features. In contrast to partial dependence plots from the Random Forest Classifier, these plots do not take confounding and interaction between features into account. Feature definition and selection were aimed at reducing differences between partial and marginal dependence, resulting in dependence plots that are independent of the model and feature selection, but still representative of the relations that the model has learnt.

## RESULTS

To illustrate model performance for a variety of species groups and different types of observations, results are shown in detail for four of the 150 species: a coastal bird species (the Eurasian oystercatcher *Haematopus ostralegus*), the most observed plant in the dataset (the ribwort plantain *Plantago lanceolata*), a moth that is mainly caught using special traps (the bright-line brown-eye *Lacanobia oleracea*) and a common mushroom (the shaggy ink cap *Coprinus comatus*).

A total of 2179 areas of one squared kilometer had at least 50 observations between 2020 and 2022 and were selected for testing the model. The number of observations in these areas varied between 50 and 38164 and had a median of 281.

For each of the 150 species, areas for testing the model were selected based on occupancy, defined as an observation in at least two out of three years from 2020 to 2022 (see red squares in Fig. 2). One species, the Asian Hornet, was filtered out because not one area met the occupancy definition. For the remaining 149 species, the number of occupied areas varied between 16 and 1561 and had a median of 147. An observation during 2023 took place if any of the visits to the area resulted in detection of the species. Among occupied areas, the percentage of observations per species in 2023 ranged from 11% for the big sheath mushroom *Volvopluteus gloiocephalus* to 86% for the hornet mimic hoverfly *Volucella zonaria* (median 59%). Outside occupied areas, this percentage ranged from 0.4% for the fly agaric *Amanita muscaria* to 35% for the common buzzard *Buteo buteo* (median 8%). Observations thus had a higher probability inside previously occupied areas, implying a correlation between occupation based on 2020 to 2022 and observation in 2023.

**Figure 2.**
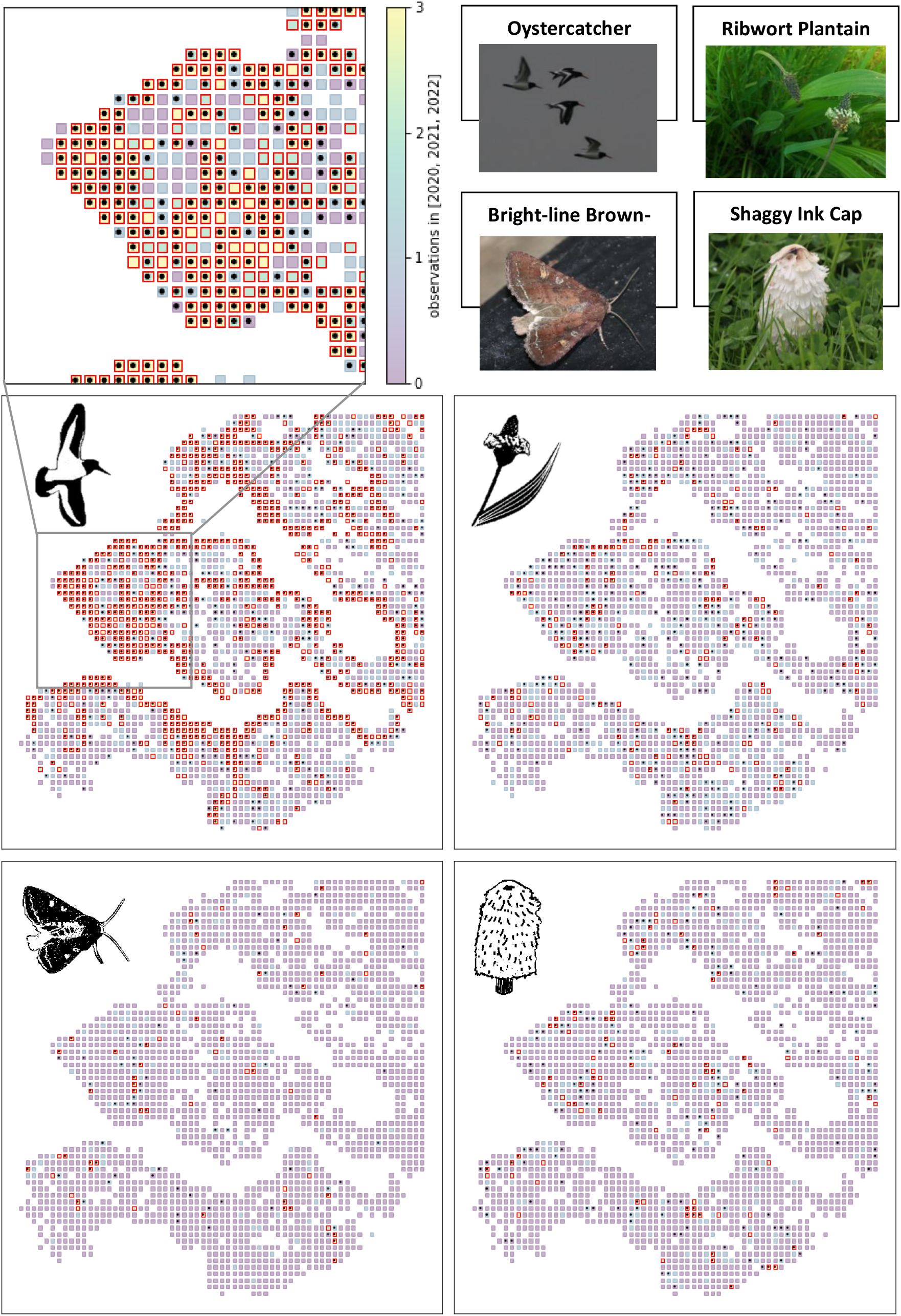
Selected areas for the modeling dataset (red squares) and observations in 2023 (black dots) for the selected species. Background color of the areas represents the number of years from 2020 to 2022 that the species was observed. Only areas where the species was observed in at least two out of three years are selected for modeling.

A Random Forest Classifier was trained per species, to learn the relation between features of each visit and observations in 2020, 2021 and 2022. The dependence plots offer intriguing insights into the factors influencing detection and, consequently, shed light on underlying sources of bias. List length is the strongest feature individually and improves the average AUC of the final model by 0.037. For all species, the detection probability increases with the list length (first column in Fig. 3). This makes sense as a more comprehensive effort during the visit leads to a larger probability that the focal species is detected and reported. The detection probability of bird and moth species increases faster with list length, which is consistent with the fact that the numbers of common birds and moths were smaller than that of plants and mushrooms, resulting in larger proportions per species. The oystercatcher and the bright-line brown-eye had even steeper curves than the average bird and moth, because both were among the most observed species. Interestingly, the shaggy ink cap shows a flatter curve that even seems to decrease when the list length increases. This may be related to the ecology of the shaggy ink cap: it grows in grasslands that are typically not very species-rich. List length tended to be largest for moth species, probably as a consequence of the strategy to capture moths using a light trap that detects all species at a single location.

**Figure 3.**
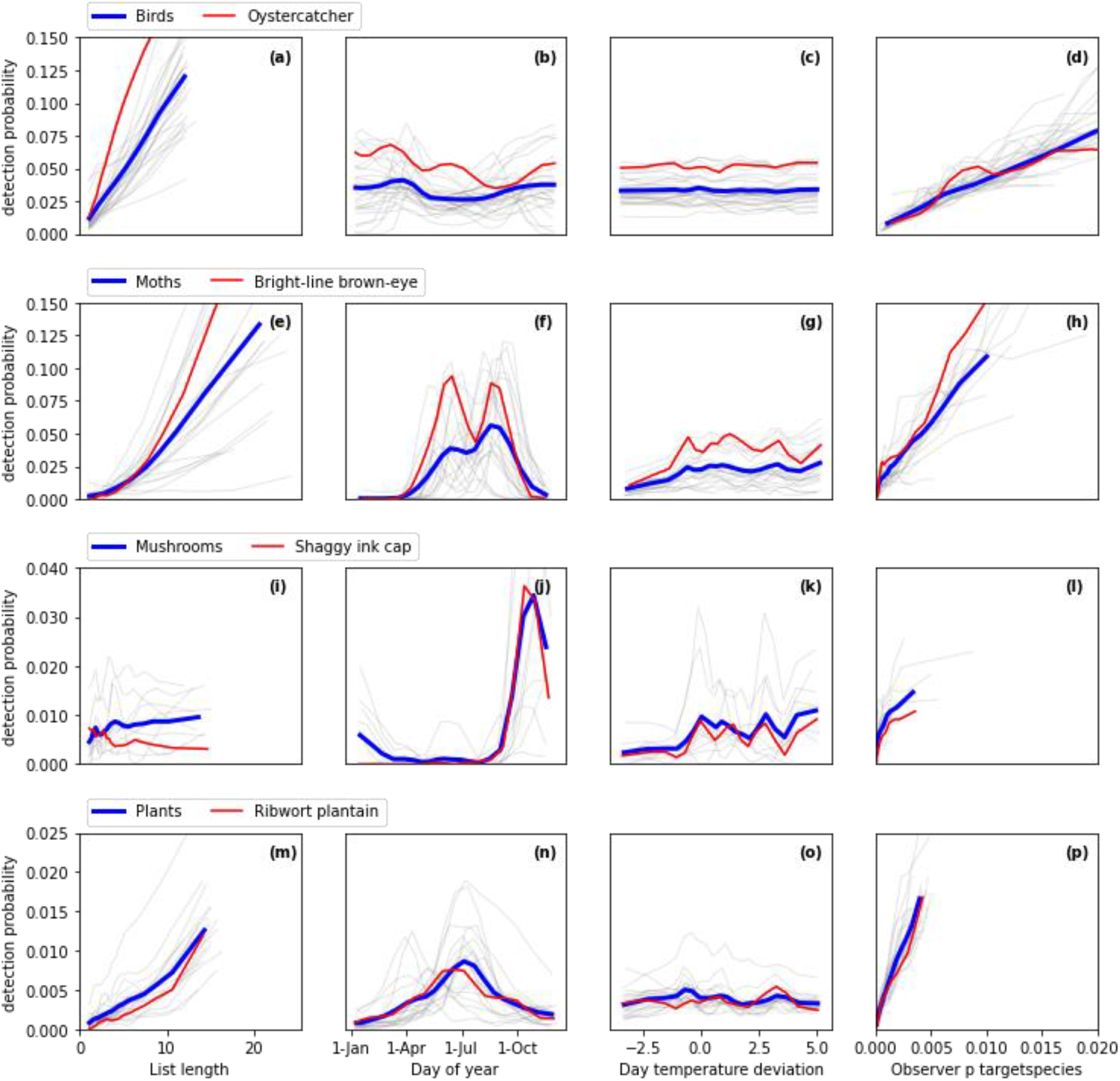
Marginal dependence plots, showing the conditional probability of detection given occupancy per species for each visit, as a function of the selected features, per species group. In grey, the individual curves of all species. In blue, the average curve over all species. In red, the four species that are covered in more detail throughout the results.

Each species group had its own typical distribution of detection probability over the year (second column in Fig. 3). The “day of year” feature strongly contributes to the final model by improving the average AUC by 0.020. Mushrooms had the highest probability of detection during visits in autumn, while moths and plants had their peak during summer. On average, birds had more or less constant detection probabilities throughout the year. Individual bird species do show curves as expected, dependent on the period that the bird usually stays in the province. Migratory birds like the barn swallow *Hironda rustica*, the western marsh harrier *Circus aeruginosus* and the common chiffchaff *Phylloscopus collybita* have their peaks during the warm season, while most other birds have higher detection probabilities during winter. Like most moths, the bright-line brown-eye is most often detected during the summer, and even two separate generations (Vajgand, 2009; Vlinderstichting, 2024) are reflected in the detection probabilities. Some plants had their peak in spring instead of in summer (see grey lines in Fig. 3n), most notably cow parsley *Anthriscus sylvestris*, ground-ivy *Glechoma hederacea* and the red dead-nettle *Lamium purpureum*, this seems to be related to the month when these plants (first) bloom.

The effect of temperature deviations in the third column of Fig. 3 was generally weak. This feature improved the average AUC of the final model by only 0.004 and was only selected because it is the strongest one compared to other weather features. However, for moths there is a trend with increased detection probabilities when temperatures were higher than the 30-year mean for that time of the year. To a lesser extent, this also seemed to be the case for mushrooms. Note that the temperature deviation could also determine how many visits took place. Our analysis does not look into that because detection probability is calculated only on the occasions when a visit did take place.

All species tend to have a higher detection probability when the observer had detected and reported the focal species often during visits in earlier years (fourth column in Fig. 3). This feature assists the model to distinguish between observers with a different focus, as some observers only report birds while others are mainly focused on butterflies. Among all observer features, the selected feature about detecting the same species is individually the strongest predictor. It is also the second most important feature in the final model, improving the average AUC by 0.034. Compared to other observer features, it shows the most consistent and interpretable dependence plots, in the sense that it shows about the same pattern for all species. Other features about the observer history were created and tested in addition to the final model (see Appendix 2), but none of the added features improved the average AUC by more than 0.01.

### Validation

The AUC has been calculated for non-detection per visit, by testing predicted non-detection probabilities against actual non-detection during the visit. Over all species, this AUC per visit varies between 0.47 and 0.97 and has a median of 0.77. These outcomes can be compared to an AUC of 0.5 that would reflect a random probability per visit, so they prove that the model is quite successful in predicting visits that result in a detection.

The non-detection probability of an area (conditional on being occupied in previous years) is the product of the conditional non-detection probabilities of all visits to the area in 2023. Over all species, the resulting AUC per area varied between 0.48 and 0.96 and had a median of 0.77. These results are compared to two benchmarks that are simpler because they do not use all available information. The first benchmark takes only the number of visits into account. For this benchmark, areas are ordered only by the number of visits, expecting a larger detection probability in areas that were visited more often. This benchmark results in a median AUC of 0.69. The second benchmark applies the same modeling approach, but only uses the list length feature, disregarding information about weather, observer and season. This more competitive benchmark results in a median AUC of 0.73. Instead of looking at the median AUC over all species, results were also compared within each species group separately. Figure 4 shows that the predictions of the full model consistently outperform both benchmarks for all the species groups, although the difference is less clear for Diptera.

**Figure 4.**
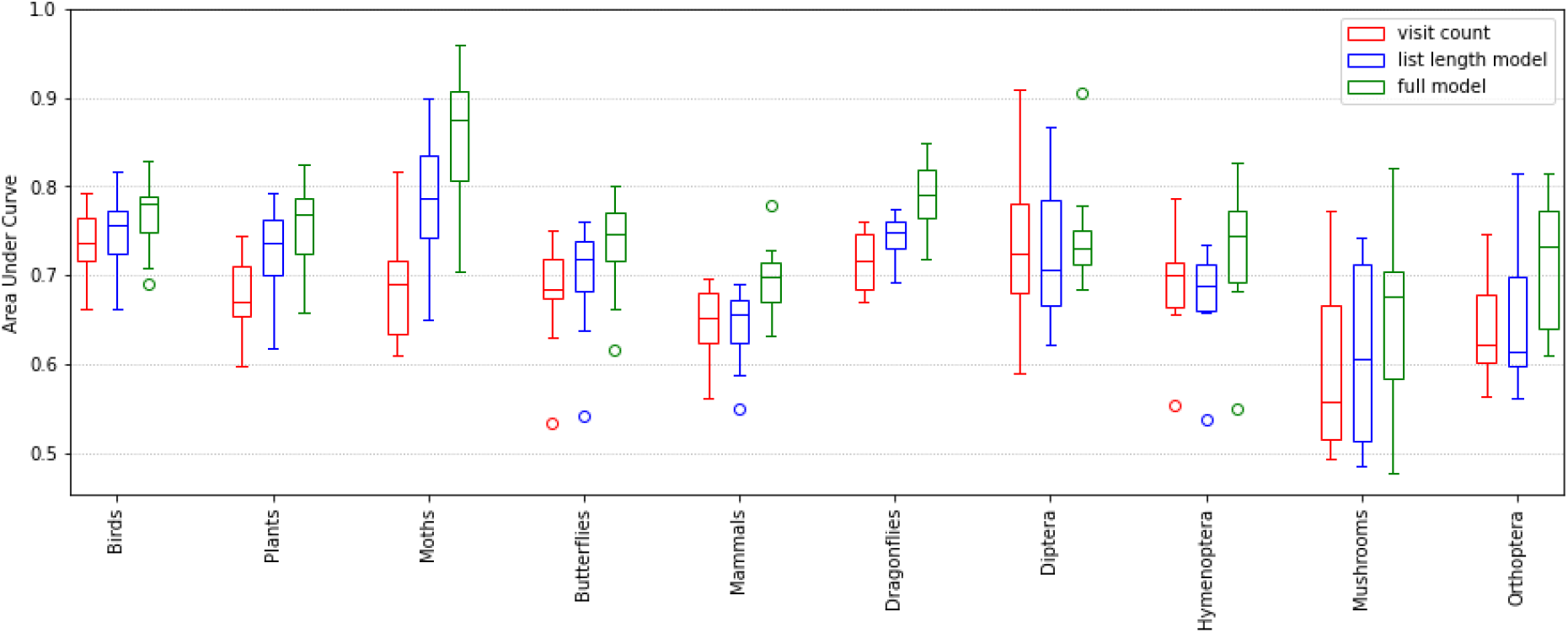
Boxplots summarizing the Area Under Curve (AUC) per species group, given three different approaches to rank the areas. Each box contains the middle 50% between the first and third quartile of the AUC outcomes for an approach carried out on a species group (for example, the red box for the visit count approach applied to birds on the left). The horizontal line in the middle of the box is the median, whiskers indicate the full range between minimum and maximum, with the exception of outliers, indicated with circle markers outside this range. The AUC is expected to vary between 0.5 for a worthless model and 1.0 for a perfect model. The figure shows that the full model beats both benchmark approaches within each species group.

The non-detection probabilities were averaged per species to compare them to the proportion of areas where the species was observed at least once in the test year 2023. The average was calculated for occupied (at least 2 years in 2020 to 2022) and ‘unoccupied’ areas separately. As expected, inside occupied areas there is a strong negative relation between non-detection probability and observations (blue dots in Figure 5). The dashed grey line shows the ideal relation for a perfect model. The results were scattered around it, but the observation percentage is a bit lower than expected when the non-detection probability gets lower and the variance seems to increase towards non-detection probabilities around 50%. In areas that were not occupied, the observation percentage was generally lower and not as strongly related to the non-detection probability: the reason for not observing a species is (at least partly) that the area was not occupied in 2023.

**Figure 5.**
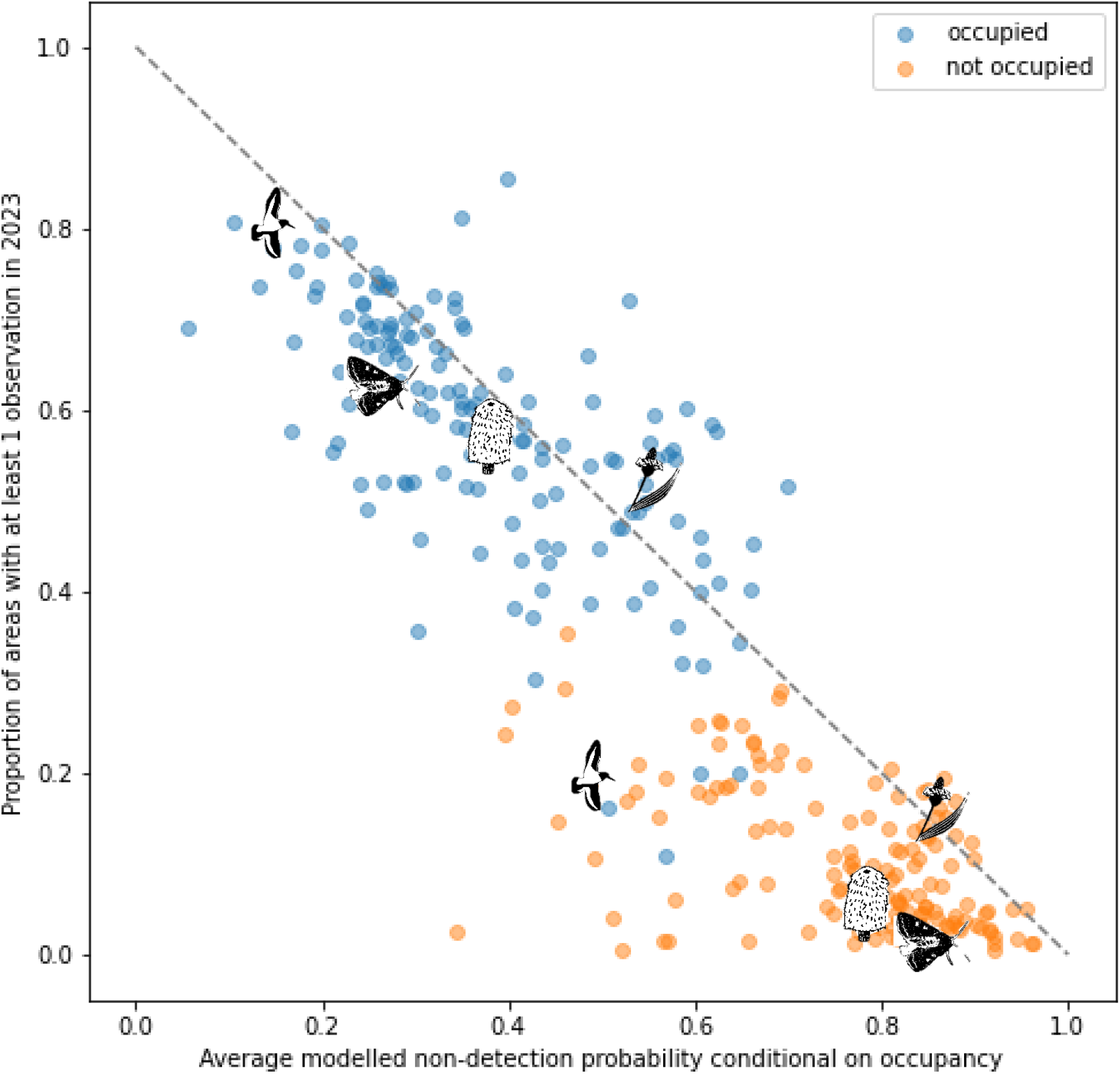
The relation between the average non-detection probability per species conditional on occupancy in previous years (horizontal axis) and the proportion of areas with an observation in the test year 2023 (vertical axis). The blue dots represent species-specific averages for areas occupied in at least 2 years in the 2020-2022 period. Model predictions for conditional non-detection probabilities were also calculated for the other (previously ‘unoccupied’) areas, which had fewer observations in 2023 and showed a weaker relationship with the proportion of observations (orange dots). In ‘unoccupied areas’ (orange dots) the ribwort plantain has relatively high proportion of observations, suggesting that a lack of detection may be the sole reason for the unoccupied status. Both the bright-line brown-eye and the shaggy ink cap show similar (high) non-detection probabilities, but a lower proportion of observations, meaning that the small effort led to even fewer observations, suggesting that the occupation status has a correlation with true occupancy. The same applies to the Eurasian oystercatcher, in a more conclusive manner because even in unoccupied areas there has been a considerable effort, given the non-detection probability around 50%, resulting in only 20% observations. In occupied areas, all four species have a proportion of observations that is in line with the complement of the non-detection probability, as expected.

The probability of non-detection (conditional on occupancy) that the model reveals, quantifies the sampling effort. In unoccupied areas, this probability is not the actual probability to not detect a species, but only the probability that a species would not be detected if it were present, given the sampling effort. Therefore, the probability of detection is a suitable quantification of (the lack of) sampling effort, both for occupied and unoccupied areas. Darker areas in figure 6 represent lower non-detection probabilities, and had a more complete sampling effort. Given all visits by citizen scientists in 2023, the sampling effort for oystercatchers turned out to be the most comprehensive one. The spatial distribution of the sampling effort looks about the same for these four species. As expected, the effort is clustered in space and highest in densely populated areas like the cities of Goes and Middelburg, and areas with a high biodiversity like areas along the coast and nature reserves that attract many observers.

**Figure 6.**
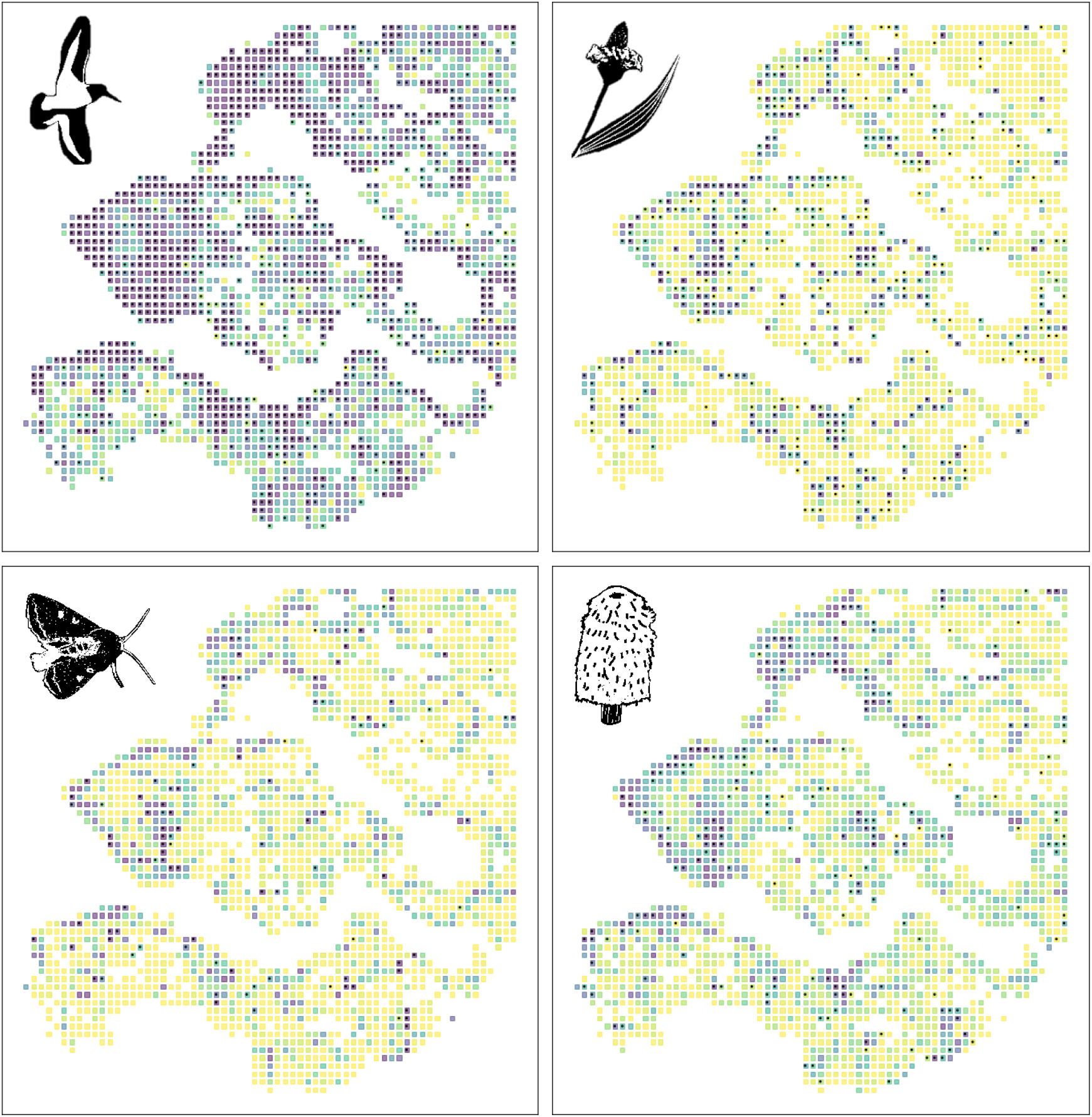
Per squared kilometer area, the probability of non-detection given occupancy, with observations in 2023 as black dots. All areas are shown, because even in (previously) unoccupied areas this probability quantifies the sampling effort. The brightest (yellow) color represents 100% non-detection probability, all four selected species have non-detection probabilities ranging from 0.00% to 100% with an average of 35.8% for the Eurasian oystercatcher, 82.9% for the ribwort plantain, 80.9% for the bright-line brown-eye and 74.3% for the shaggy ink cap.

## DISCUSSION

### Key results

Citizen science data about biodiversity are a growing source of information that is often left unused because it is hard to derive representative insights from observations that do not follow a known protocol. The fact that only presence data are recorded is challenging because it is unknown what lead the observer to select some and not other species for reporting. The method proposed in this paper gives insights into causes of bias, the non-linear effects of these causes and their relative importance. Based on input features about the visit, the season, the weather conditions and the observer, a model predicts the probability of non-detection when the area is occupied by the species of interest. Validation results confirm that the model succeeds in predicting the non-detection probability based on the available features.

The model and its predictions focus on detection only, because the model has been fit on a subset of areas that are known to be occupied by the species. Therefore, occupancy does not play a role in the relations that the model finds. This implies that the predictions are suitable to quantify sampling effort. This effort is not just based on the number of visits, but also takes circumstances during the visit that have an impact on detection probability into account.

By fitting models for detection inside occupied areas only, complexity has been considerably reduced. In contrast to other studies, we did not need to limit the number of covariates, assume linear relationships, fix detection probabilities per species (Kéry and Royle, 2008) or per visit (Outhwaite et al., 2018) or filter species that show low detection probabilities (Ferrer-Paris & Sánchez-Mercado, 2020). Moreover, by using external data to define occupancy, we avoided the reliance on a strict definition of occupancy to estimate the detection probability from replicate samples.

### Selection of occupied areas

Occupied areas need to be selected in order to fit the model. It is required that occupancy is static during the time period of the modeling dataset. This selection can be challenging for species that are fluctuating, expanding or declining in range. Because of this challenge, occupied areas for the rapidly expanding Asian hornet could not be established based on historical records.

The modeling approach allows a very loose definition of occupancy. Occupancy does not need to mean that a species is present at the time of the visit, like regularly assumed (Efford & Dawson, 2012). When a bird of prey went hunting somewhere else, a butterfly is unfindable because it is a pupa or a migratory bird is spending the winter in Africa, their area would still count as occupied. Given the features of the visit, the model will soon recognize that the detection probability is lower or zero during a certain season or weather condition. Within occupied areas, all complexity in patterns of species presence is taken care of by the model. The model could even find interactions between species behavior and observer habits, for example an observer who is more likely to detect a migratory bird when it first returns to its breeding area. A different type of occupation among areas within the study area is challenging, however. For instance, many oystercatchers occupy the tidal areas in winter, while in the breeding season they are spread out from the coast to far inland (Allen et al., 2019). A pattern like that can confuse the model, because all areas share the same parameters and seasonal patterns are assumed to be the same among areas.

Selection of areas that are actually not occupied may introduce bias in the modeling results. The dataset would then contain visits with non-detection that are not caused by detection, but by (the lack of) occupancy. That would lead to overestimation of the non-detection probabilities. Treating unoccupied areas as occupied areas could be a consequence of misidentifications (Miller et al., 2011), but otherwise it is not likely given our approach: the relaxed definition of occupancy includes occasional visitors, and could be made more stringent with the definition of ‘previously occupied’, e.g. reported in all three previous years, or multiple times per year. The opposite, missing occupied areas for the selection, would not need to be a problem. The only assumption required to have realistic results is that the areas in the dataset are representative of all the occupied areas.

In this study, the selection of occupied areas is a consequence of detection in earlier years. When the distribution of sampling effort over areas is very skewed and persistent over years, the representativity assumption may be violated. Moths, for example, are often caught using light traps that are only applied at certain sampling locations for years in a row. Within the selected areas that are occupied by moths, the detection probability is quite high. In other areas that are not part of the modeling dataset, the detection probability is much lower and will be overestimated by the model that is mostly trained using areas with light traps. The challenge is to add features about observer or area that will help the model to recognize the difference between areas with and without light traps.

The current feature about the percent of target species in the observer history does help to partly relieve this problem, because it distinguishes visits of observers who previously reported the same moth species – these are the observers involved with light traps and likely to find and report moths even when they become active in new areas. Attempts to define features at the area level, like the percent of observations in the species group of the focal species, did not improve the results and introduced the challenge to avoid that the model learns to distinguish between areas instead of area characteristics.

Another option to select occupied areas would be to use an external dataset that functions as the gold standard. For example, in parts of this study area, shorebirds are systematically counted multiple times a year by professionals following a standardized protocol (Lilipaly & Sluijter, 2023). Using these data to define occupancy by shorebird species would lower the risk of systematically selecting more areas as occupied when they have a higher sampling effort that is persistent over years, because the protocol makes sure that the sampling effort is kept constant. An additional advantage is that occupancy could be established during the same time period, instead of the history within the same dataset. But there are also additional challenges when using an external dataset: the resolution in time and space may not match well and there may be possible overlap between datasets because observers submit their observations to multiple platforms.

When observations in the test year 2023 were compared between occupied and previously unoccupied areas, it turned out that species were more often found in the same areas where they were previously found. This correlation can be caused by occupancy itself: a previously occupied area has a larger probability of being occupied again. But an alternative explanation might be the sampling effort: areas that are consistently visited more often have a higher probability of observation, both in 2020 to 2022 and in 2023. Figure 5 shows that both things may play a role and that the model helps to distinguish between the two. Regarding the sampling effort, the average non-detection probability tends to be higher in unoccupied areas: this confirms that some of the unoccupied areas are not well investigated (with regard to the focal species) in 2023, which makes it more likely that their unoccupied status based on 2020 to 2022 has been a matter of sampling effort. On the other hand, there is a clear distinction between occupied and previously unoccupied areas that have the same average non-detection probability: given the same sampling effort, observations in 2023 are more likely in areas that were occupied in at least two years in the 2020-2022 period.

### Extensions

All calculations and models applied in this paper are species-specific. Therefore, attributes of species are constant values and not suitable as features for modeling. Some attributes like size, visibility, recognizability, species group and rarity are likely to have an impact on detection. In the current approach, this will affect only the average level of predictions per species. An interesting extension would be a meta-model to describe and predict how attributes of species influence the detection probability. This would be particularly useful to create predictions for species that have very few records of observations.

The modeling of non-detection facilitates improved information about the distribution of species, by estimating and correcting how various features cause bias. The steep increase in the number of observers that participate in citizen science makes it even more challenging to estimate trends over time (Fink et al., 2023). In principle, the model could recognize changes in detection probability as a consequence of more observers and observers with different characteristics. It would require additional feature extraction and validation to extend the model to apply it for estimation of trends.

In practice, abundance of species will have an impact on the detection probability. In the dataset for this study, abundance estimates are not standardized and often missing. Therefore, species presence has been simplified to occupancy only, disregarding abundance. It would be interesting to extend the model to take abundance into account as a feature that influences the detection probability. This may be a complicated puzzle because occupancy is also used to select areas for modeling. Incomplete information in existing records is not uncommon in citizen science data. Other potentially interesting features that have not been used because of incomplete data include time of day and life stage of the observed specimen.

This study does not look into the causes of relations between visit features and detection probability. The model learns the relations between input features and detection to produce a prediction, without necessarily understanding what the relations mean. Some of the relations may be based on artefacts in the data that do not actually help to correct bias. Better understanding of the relations will help to create more accurate models. This requires both expert insights and investigation of how the model creates its predictions.

Finally, our detection model only focuses on non-detection bias, without considering misidentification that could lead to false positives. Miller et al (2011) show that uncertain detections can be used to improve occupancy estimates. For the approach in this study, uncertain detections based on indirect observation (scat or tracks) would not be problematic because the definition of occupancy does not require instantaneous occupancy. Uncertain detections because of unskilled citizen scientists would usually be detected by validators who check feasibility and evidence in the form of photos that are uploaded.

## FURTHER RESEARCH

The bias in citizen science data is a consequence of the specific interests and habits of observers. For direct inference about occupancy, the entangled causes of occupancy and detection need to be unraveled. To achieve that in the context of citizen science data, both limitations of model complexity and additional assumptions would be required. By focusing on detection within occupied areas, we deliberately took an observation-driven view (Royle & Dorazio, 2008). The consequence is that our model does not allow for direct inference about occupancy. The quantification of sampling effort needs to be combined with other information to arrive at conclusions about occupancy, the quantity of interest. For many species, this kind of information is available in datasets that do contain systematically gathered data, but only in a relatively small subset of areas. Koshkina et al. (2017) prove that a considerable improvement can be attained by combining citizen science and systematically gathered data. They combine both sources in an integrated model, that is quite complex and has issues to correctly identify different sets of covariates. Following the approach presented in this paper, it would be quite natural to have separate models to exploit the information from these sources before it is combined.

### Combining conditional models

The conditional model for detection purposely neglected occupancy by focusing on occupied areas only. To model suitability of the areas for occupancy, we suggest fitting separate SDMs (Elith & Leathwick, 2009) using explanatory variables that represent the physical environment and the local climate. Ideally, these models are fitted using some set of areas where complete presence and absence data are available, in a way that the assumption of perfect detection makes sense. This corresponds to the model displayed in the left part of Figure 1. In contrast, the detection models discussed in this paper only use visit specific explanatory variables like weather, observer and season. Although some authors argue and prove that the physical environment does have an impact on detection (Gu & Swihart, 2004; Chen et al., 2009) there is a risk that these variables would mainly be related to abundance as a cause of detection. For example, think of a model that uses the number of trees in an area as an input to predict detection of a bird species that lives in trees. It is likely that more trees would provide more hiding places for the birds, making detection harder – which would be a reason to include the number of trees in the model. But it is also quite likely that the number of trees would change the abundance of the tree-living bird species in that area. The model simplifies presence of the bird to occupancy, so this increase in abundance would not change the representation of the bird in the data. But abundance of the bird might very well change the likelihood of detection, introducing a relation between environmental characteristics and detection that is mediated by abundance and opposite in direction to the direct effect of trees on detection probability. Including environmental characteristics to predict detection might improve predictions of detection probability, but it contradicts both the choice to estimate occupancy instead of abundance and the choice to build separate conditional models for detection and occupancy.

### Correcting bias

A promising application of the conditional model presented in this paper is the potential to correct bias in citizen science data. To do so, the SDMs in the previous paragraph need to be combined to the detection models we presented. First, suitability-based probabilities of occupancy can be estimated for any area, based on SDMs that take climate and environment into account. Next, citizen science data can be used to update these expectations. In areas without presence records, non-detection probabilities according to the detection model provide information about the sampling effort, based on visit specific explanatory variables. Non-detection inside an occupied area is quite likely if the sampling effort has been small, but becomes less likely with an increasing sampling effort. This gives an excellent opportunity to combine the information that is available in systematically gathered data on one hand and citizen science on the other hand. Probabilities of occupancy can be estimated for all areas by applying the SDMs that were learnt in a subset of areas where complete presence absence data are available from systematic research. These estimates are based on environmental characteristics and therefore represent the suitability of the area for occupancy. To estimate the distribution of a species, this suitability needs to be translated into an occupancy score. This could be done using a fixed threshold (Liu et al., 2005) but it would be more sophisticated to take into account how well the area has been investigated. If the sampling effort has been very substantial in an area without detecting the species, it is less likely that the area is occupied, even when it is very suitable given the environmental characteristics. That is where citizen science data can be combined to improve the estimates for areas outside the systematically monitored areas. The sampling effort, quantified by taking the circumstances during all visits into account, tells how well the area has been investigated – and how likely it is that the area is occupied when a species has not been reported after this effort took place. To summarize this, the suitability-based probability of occupancy needs to be adjusted downward when citizen science data do not contain presence records while there has been a substantial sampling effort.

This approach to correct bias in occupancy estimates based on citizen science can be formalized using a mathematical theorem known as Bayes’ rule (Bayes, 1763). To estimate the posterior probability of occupancy in areas where presence has not been ascertained, this rule updates the suitability-based prior probability of occupancy using the conditional probability of non-detection given occupancy. The resulting posterior probability of occupancy given non-detection is suitable to draw conclusions about occupancy in areas where the species has not been detected. The posterior probabilities can be validated by comparing them to external data about occupancy.

## Appendix 1.

Illustration of how the modeling dataset was created, using the preceding years to determine occupancy per area for each year separately. For example, to qualify an area as occupied by a species in the year 2020, observations in at least two years of 2017, 2018 and 2019 were required. Training set visits to occupied areas in 2020, 2021 and 2022 were used for training the model. Test set visits to occupied areas in the year 2023 were used for testing the model, by calculating the AUC metric.

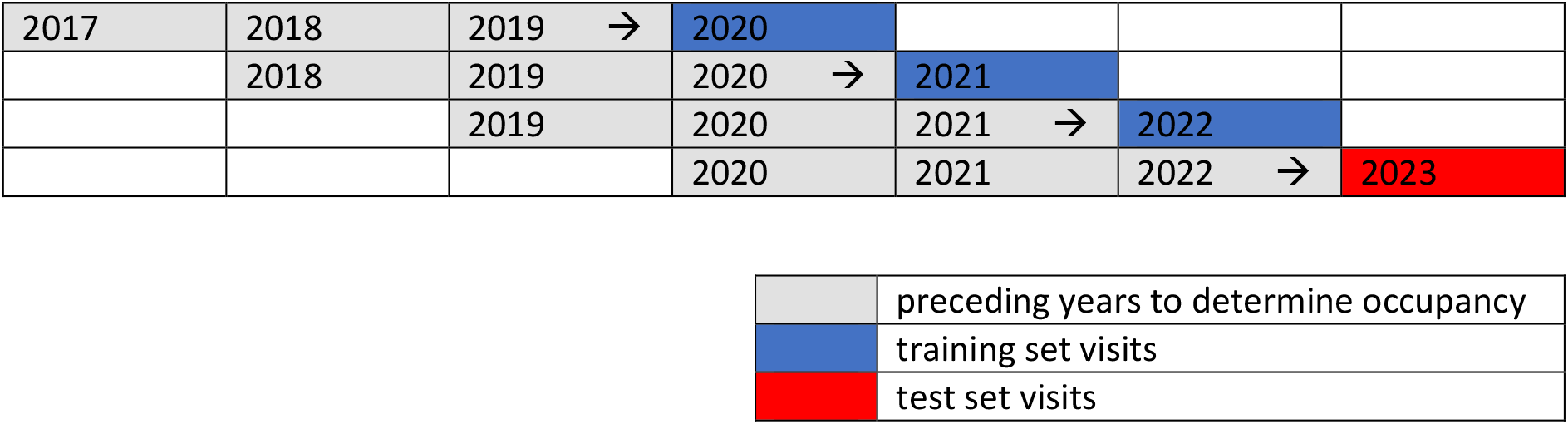

|  |  |  |  |  |  |  |
| --- | --- | --- | --- | --- | --- | --- |
| 2017 | 2018 | 2019 → | 2020 |  |  |  |
|  | 2018 | 2019 | 2020 → | 2021 |  |  |
|  |  | 2019 | 2020 | 2021 → | 2022 |  |
|  |  |  | 2020 | 2021 | 2022 → | 2023 |

|  |  |
| --- | --- |
|  | preceding years to determine occupancy |
|  | training set visits |
|  | test set visits |

## Appendix 2.

Additional features that were created but not selected, listing the improvement of the average AUC when only that feature was added to the final model.

| Feature | Average AUC improvement, when added to the final model that includes the 4 selected features |
| --- | --- |
| Total number of species, reported by the observer before 2023 | 0.005 |
| Average number of observations per species, reported by the observer before 2023 | 0.004 |
| Mean species rarity of observations, reported by the observer before 2023 | 0.001 |
| Mm precipitation on the day of the visit (transformed using the natural logarithm of mm+1) | 0.001 |

## Appendix 3.

Marginal dependence plots, showing the conditional probability of detection given occupancy per species for each visit, as a function of the additional features that were created but not selected, per species group. In grey, the individual curves of all species. In blue, the average curve over all species. In red, the four species that are covered in more detail throughout the results.

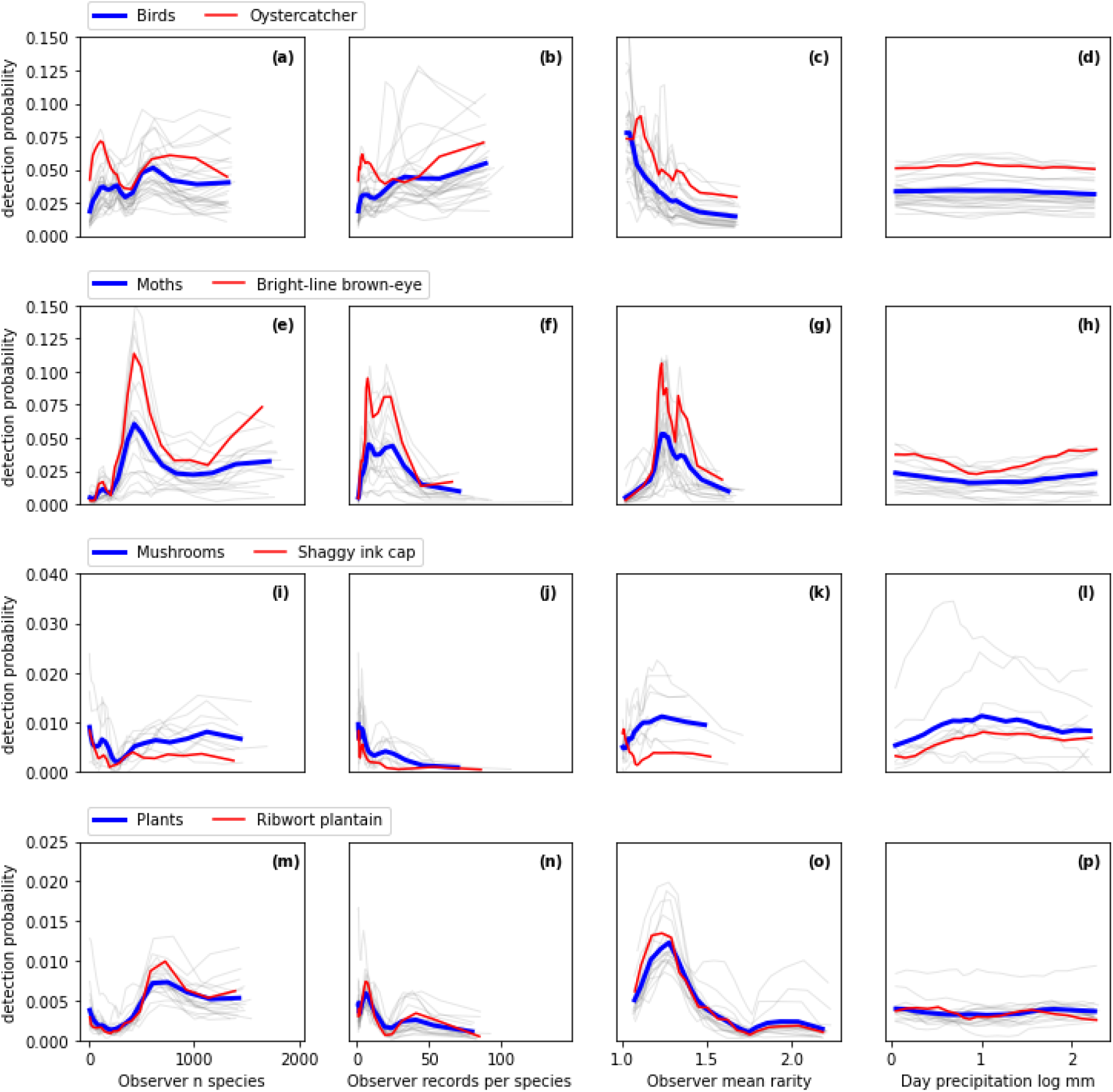

## Footnotes

1 Based on unique observation IDs; during one excursion many observations can be recorded, even of the same species within the same area

